# Likelihood-based signal and noise analysis for docking of models into cryo-EM maps

**DOI:** 10.1101/2022.12.20.521171

**Authors:** Randy J. Read, Claudia Millán, Airlie J. McCoy, Thomas C. Terwilliger

**Affiliations:** Department of Haematology, Cambridge Institute for Medical Research, University of Cambridge, Hills Road, Cambridge CB2 0XY, United Kingdom; SciBite Limited, BioData Innovation Centre, Wellcome Genome Campus, Hinxton, Cambridge CB10 1DR, United Kingdom; New Mexico Consortium, Los Alamos National Laboratory, 100 Entrada Drive, Los Alamos, NM 87544, USA

**Keywords:** Likelihood, cryo-EM, docking, information gain

## Abstract

Fast, reliable docking of models into cryo-EM maps requires understanding of the errors in the maps and the models. Likelihood-based approaches to errors have proven to be powerful and adaptable in experimental structural biology, finding applications in both crystallography and cryo-EM. Indeed, previous crystallographic work on the errors in structural models is directly applicable to likelihood targets in cryo-EM. Likelihood targets in Fourier space are derived here to characterise, based on the comparison of half-maps, the direction- and resolution-dependent variation in the strength of both signal and noise in the data. Because the signal depends on local features, the signal and noise are analysed in local regions of the cryo-EM reconstruction. The likelihood analysis extends to prediction of the signal that will be achieved in any docking calculation for a model of specified quality and completeness. A related calculation generalises a previous measure of the information gained by making the cryo-EM reconstruction.

## 1. Introduction

The problem of docking models into cryo-EM maps is similar to the molecular replacement (MR) problem in crystallography. The key difference is that cryo-EM data are enriched by the phase information that is lost in crystallography, and the resulting increase in signal-to-noise greatly simplifies the task of translating an oriented model with FFT-based correlation functions. However, this phase information cannot be used directly in assessing different model orientations prior to the translation search, so many existing docking algorithms rely on a systematic six-dimensional search over possible orientations and positions.

In crystallography, MR algorithms have been made significantly more sensitive by using likelihood scores for rotation, translation and rigid-body refinement tasks (McCoy *et al*., 2007). In addition, an understanding of the relationship connecting data and model quality with the likelihood scores that can be expected in a particular calculation has opened up new possibilities for tailoring the MR calculations to the problem at hand (McCoy *et al*., 2017; Oeffner *et al*., 2018). These concepts can be applied to the related problems in cryo-EM.

Most existing docking methods for cryo-EM are scored by variants of cross-correlation functions. Comprehensive reviews of these score functions have been compiled by others: (Zundert *et al*., 2015; Cragnolini *et al*., 2021). Some examples include cross-correlation of the experimental cryo-EM map and a map computed from coordinates (Stewart *et al*., 1993), local cross-correlation (Roseman, 2000), Laplacian filtered cross-correlation (Wriggers, 2012) and core-weighted cross-correlation (Wu *et al*., 2003).

When comparing a variety of scores of fit to model, including cross-correlations, Joseph *et al*. (2017) found that mutual information was a better discriminator for low to medium resolution maps. Like the likelihood score proposed here, mutual information is a probabilistic measure, but it works with real-space voxel values, not Fourier terms. In addition, mutual information does not explicitly account for errors in the reconstruction itself.

As noted below, our docking target is based on similar ideas to the likelihood-based refinement target for models against cryo-EM maps used in Refmac (Murshudov, 2016), but differs importantly in using a more sophisticated error model for experimental data that takes account of the directional dependence of both the signal and the noise in Fourier space.

## 2. Probabilities and likelihood targets

### 2.1. Error model for single-particle cryo-EM data

For a cryo-EM reconstruction, the aim is that each individual molecule or molecular assembly in a particle is essentially a rigid object, either by nature or as a result of particle selection.

Errors in cryo-EM reconstructions come from a combination of suboptimal relative orientations of individual images, structural differences and imaging limitations and artefacts among the collection of particles used in the reconstruction, reviewed for instance by Ramlaul *et al*. (2019). In the individual 2D particle images derived from a series collected over the total exposure, the images are smeared by any uncorrected sample motion, degraded by effects of any radiation damage and limited in resolution by the detector pixel size. Additional random shot noise comes from counting statistics and the presence of irreproducible features in the vitrified solvent around them.

For reconstruction, the information contained in the 2D image is converted into its Fourier transform, which comprises a 2D slice through the Fourier transform of the molecule; errors in the Fourier terms can arise, for instance, from errors in the contrast transfer function correction. If the correction terms have been optimized, their values and errors will differ in different images, so we can expect the remaining errors in data from these individual images to be largely uncorrelated with particle orientation or with the images themselves. Nonetheless, if systematic errors remained it would be difficult to distinguish them from signal.

Each particle imaged in a data set will be in a different orientation and (to a greater or lesser extent) a different conformation. 3D classification will allow significantly different conformations to be grouped together, but variation will remain within the groups, corresponding in real space to blurring of the atoms over their range of possible relative positions when constructing a 3D image. Further blurring will come from uncertainties in the orientation and position of the particle in each image, when averaging the Fourier terms from different images to obtain a 3D data set.

In our error model, we consider that the signal in an individual Fourier term in the reconstruction comes from the Fourier transform of the image of atoms at rest, blurred by the effects of global and local variations in orientation and position. These blurring effects are similar to what is modeled locally in crystallography by anisotropic displacements or, on a larger scale, by translation-libration-screw (TLS) models (Schomaker & Trueblood, 1968).

Random noise in contributions from individual particle images will be reduced when corresponding Fourier terms from different images are averaged. However, the existence of preferred orientations will mean that the magnitude of the random noise terms, after averaging over different numbers of observations, will vary with direction in Fourier space. In principle, this could be modeled by keeping track of redundancy during the reconstruction process, but in the current implementation we are starting from conventional half-maps rather than individual particle images. We assume that the particles chosen for the half-map reconstructions are chosen randomly, so that the half-maps contain different random selections from the same sources of error. We approximate the directional and resolution dependence as a smoothly-varying function in Fourier space. Since variations in conformation need not be correlated with orientation preference in the sample, the two sources of variation in signal and noise are evaluated independently.

If we consider smaller sub-volumes of the full reconstruction (useful when searching for small components, as discussed below and in the accompanying paper (Millan *et al*., 2023)), the strength of the signal will vary because the degree of local structural order varies. On the other hand, the strength of the noise should be reasonably uniform over different parts of the full reconstruction, as discussed by Palmer and Aylett (2022). In the algorithms described below, we nonetheless determine the noise power independently for sub-volumes, because cryo-EM reconstructions are commonly masked towards the periphery of the cube, and the effects of this can vary among sub-volumes.

Because the estimation of noise requires the comparison of independent measurements, all of our signal and error evaluation is carried out using the Fourier terms computed from the halfmaps. The signal power is deduced from correlations between the half-map terms and the error power from their differences.

The signal in matching pairs of Fourier terms derived from the half maps can be expressed in terms of the underlying Fourier transform of atoms at rest (represented as **T**_*hkl*_ and drawn from a complex normal distribution with variance Σ_*T*_ representing the scattering power), multiplied by a scale factor combined with a term that varies with resolution and direction in Fourier space (represented as *A_hkl_*). The choice to separate *A_hkl_* from **T**_*hkl*_ was made to allow a Bayesian prior to be considered for the Σ_*T*_ component of the signal strength, as discussed below.

The noise term, **ε**_*hkl*_, is drawn independently for each half map from a complex normal distribution with variance Σ_*E*_. Note that both Σ_*T*_ and Σ_*E*_ will vary with resolution; as noted above, Σ_*E*_ will also vary with direction in Fourier space.

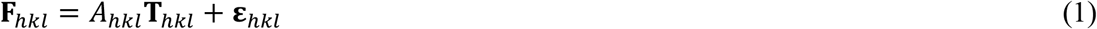

When describing the individual half-map terms, the subscripts *hkl* will be implicit for simplicity of notation:

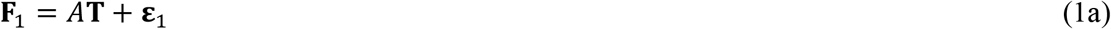

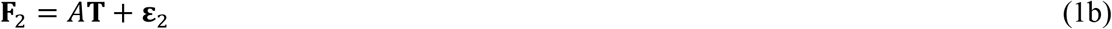

Because **T** and ***ε*** are both drawn from complex normal distributions, the joint distribution of **F**_1_, and **F**_2_ can be defined in terms of a bivariate complex normal distribution. The covariance matrix for this distribution is given by:

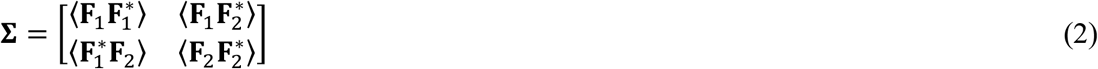

The terms in the covariance matrix can be simplified in terms of the variances of the distributions for **T** and **ε**, noting that **ε**_1_, and **ε**_2_ are independent so that their covariance is zero.

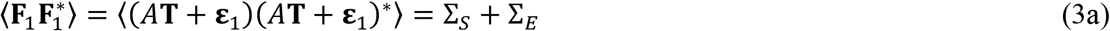

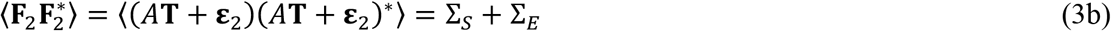

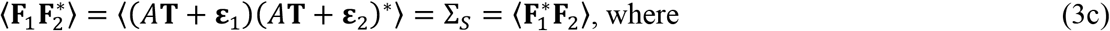

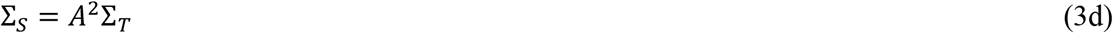

The parameters characterising the bivariate complex normal distribution can be estimated by maximising the likelihood of measuring the data derived from the two half-maps.

The determinant and the inverse of the covariance matrix are needed to compute the likelihood target:

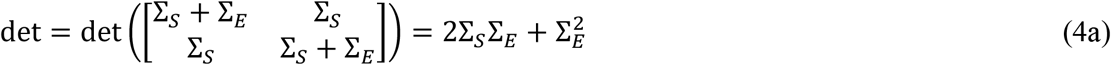

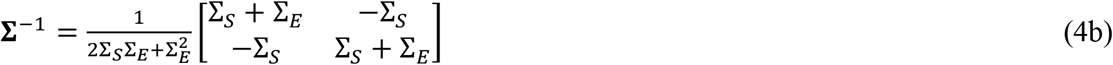

The joint probability distribution of **F**_1_, and **F**_2_ is given by the following bivariate complex normal distribution

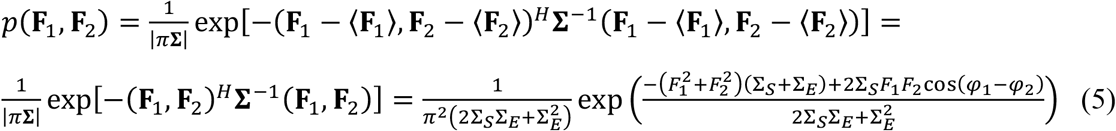

where superscript *H* indicates the Hermitian transpose. Note that, before we have any knowledge of the values of **F**_1_, and **F**_2_, their expected complex values are zero. In the final part of (5), the argument of the exponential is expanded out by multiplication, using the expressions for the inverse and determinant of the covariance matrix in (4), and the Fourier terms are represented in terms of their amplitudes and phases. The likelihood function (*L*) for a pair of half-map Fourier terms is the probability distribution in (5), given values for the variance parameters that are being evaluated. The contribution of a single Fourier term to the log-likelihood function is therefore given by the logarithm of the joint probability distribution, shown in (5).

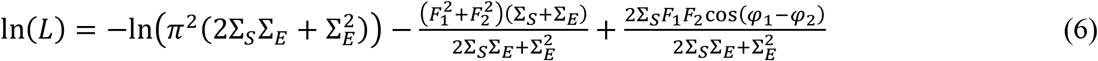

#### 2.1.1. Estimating the variance parameters in the error model

We have tested two approaches to determining values for Σ_*S*_ and Σ_*E*_ as they vary over Fourier space. One is to assume that their values are close to constant in a small region of Fourier space such as a sphere around a particular Fourier term (or at least that their variation over that sphere is such that their mean value is representative of the term at the centre). This approach makes no assumption about the functional form for their dependence on resolution or direction in Fourier space. The second approach is to assume that the variation can be captured by some combination of a resolution term (such as a constant for each spherical shell) and an anisotropic tensor representing a 3D Gaussian.

For the local variation approach, there is an analytical solution for the Σ_*S*_ and Σ_*E*_ terms that maximise the log-likelihood for a local region in Fourier space. This is obtained by taking the derivatives of the sum of the log-likelihood in (6), over a set of *n* Fourier terms, with respect to Σ_*S*_ and Σ_*E*_, then solving the simultaneous equations to find the values where the two derivatives are equal to zero.

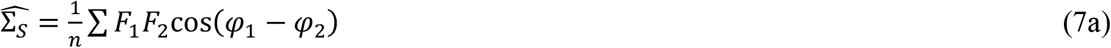

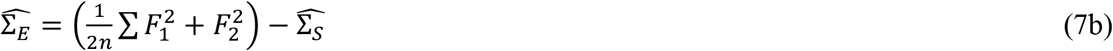

The results for the maximum likelihood estimators are intuitively reasonable. 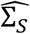 is an unnormalized correlation function of the half-map Fourier terms. If we substitute the expression for 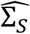 (7a) into the expression for 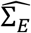 (7b), we can see that 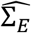 is proportional to the mean-squared value of the magnitude of the difference between the complex values of **F**_1_, and **F**_2_.

These maximum likelihood estimates of the parameters have the great advantage that they require no assumptions about the shape of their distribution over Fourier space. This is particularly acute for the Σ_*E*_ error terms; these will largely reflect the distribution of favoured orientations, which can have a number of modes that do not necessarily obey any symmetry. *A priori*, there seems little reason therefore to expect this distribution to be described well by an anisotropic tensor. Indeed, preliminary work testing the anisotropic tensor approach gave poor results (not reported here in further detail). Local variation in Fourier space is therefore used exclusively for the error terms.

There is a different trade-off for the Σ_*S*_ signal terms. In regions of Fourier space where the signal is much lower than the noise, the statistical error in the correlation function becomes very large relative to the true value, leading to serious artefacts near the resolution limit when applying the local variation approach. However, the local variation estimates of Σ_*S*_ behave well in regions of Fourier space with strong signal, with the benefit of avoiding any assumptions about the shape of the distribution of signal strength. For that reason, a hybrid method that couples the local variation approach with an anisotropic tensor approach has been chosen, as discussed below.

For the Σ_*S*_ signal terms, the assumption that the local molecular structure undergoes displacements that can be captured by an anisotropic tensor is easier to justify, based on bonding constraints, than for the Σ_*E*_ terms. An overall anisotropy model is frequently used for diffraction data in crystallography, as a relatively simple and useful approximation to accounting for overall effects of atomic displacements. For cryo-EM, such an approximation is best justified when considering a limited volume containing a component undergoing overall rigid-body displacements, but will degrade if different components undergo different rigid-body displacements.

There does not appear to be an analytical solution for maximum likelihood estimates of the terms determining Σ_*S*_, so an iterative refinement is required. The refinable parameters for the signal power in the log-likelihood function are the parameters determining the values of *A* and Σ_*T*_.

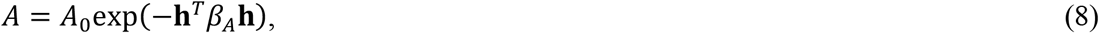

where *A*_0_ is an overall scale, **h** is a triplet of *h,k,l* values and *β_A_* is an anisotropic tensor that captures the effects of overall anisotropic displacements of the object in the map.

Σ_*T*_ is a function of resolution because the spectral variation of the Fourier transform reflects both the width of atomic features and favoured interatomic distances within the imaged object. If the signal-to-noise were reasonably high for all resolution ranges, Σ_*T*_ could be estimated reliably in resolution bins, but this is not usually a safe assumption towards the resolution limit. For a similar problem in normalising crystallographic data (Read & McCoy, 2016), we have found that a Bayesian framework using a prior probability distribution is useful: we assume that the overall spectral variation of data should be similar to the average seen in a large variety of structures, as captured by the BEST curve tabulated by Popov and Bourenkov (2003). This curve will not be exact for any particular data set, so some variation must be allowed; this is accomplished by refining bin-wise resolution parameters that are set initially to one, and weakly restraining the logarithm of their values to zero. Weaker restraints are used at low resolution than at high resolution, because the low-resolution Fourier terms depend more on molecular shape than favoured interatomic distances, and thus vary more from structure to structure. The restraints have very little effect on the refined parameters for strong data but dramatically improve the behaviour of refinement for weak data. Note that the BEST curve was derived using a large set of X-ray diffraction data to high resolution. A related curve is not yet available for cryo-EM or electron diffraction data, although a similar use of power spectra has been suggested in the past (Scheres, 2012), but we note that the spectral variation of the Fourier terms from cryo-EM reconstructions at high resolution show similar behaviour to those from X-ray diffraction, because of the predominant effect of favoured distances. Using the BEST curve, the refinable parameters for Σ_*T*_ are the resolution bin parameters in the following equation:

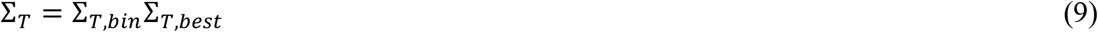

Note that the assumption that the spectral variation will follow the BEST curve will be violated when the half-maps have been manipulated, for instance by applying band-pass filters. For this reason (among others), our method requires the availability of unfiltered, unmasked half-maps.

In the hybrid method for estimating signal power, Σ_*S*_ is computed as a weighted average of the local variation estimate and estimate from the anisotropic tensor approach. Because the anisotropic tensor parameters are estimated from a likelihood target in which they contribute even when the local signal is strong, their refinement is stable, but they only dominate the determination of the hybrid estimate of Σ_*S*_ when the local signal is weak. The relative weight is determined by a sigmoid function of the local complex correlation (or local Fourier sphere correlation *CC_sphere_*) of the Fourier terms:

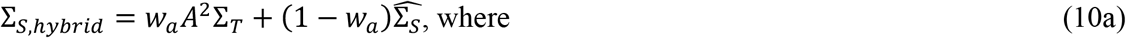

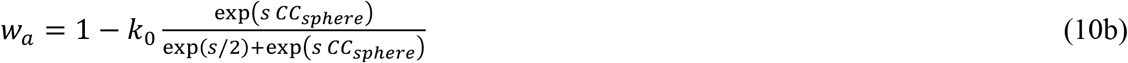

In (10b), *s* is a steepness parameter, set by default to 9, and *k*_0_ is a parameter, set by default to 0.95, to ensure that the anisotropic tensor approach still considers data with a high local correlation.

### 2.2. Likelihood target for evaluating models in cryo-EM reconstructions

To derive a likelihood target for evaluating the fit of models to data, we need to account for errors in the model (in addition to estimates of measurement error discussed above). Both structure refinement and docking can be carried out using a likelihood target that evaluates the likelihood of the map given the model. We start by considering the errors between the Fourier terms corresponding to the (unknown) true map (**T**) and the average map coefficients obtained from the two half-maps. This can be evaluated by considering the definition of the half-map Fourier coefficients in terms of the true coefficients:

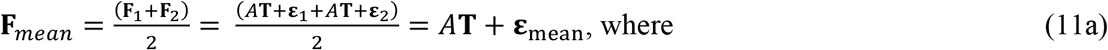

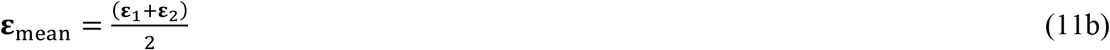

When **ε**_1_, and **ε**_2_ can be considered independent with equal variance (as we assume for halfmaps), the variance of their mean is reduced by a factor of two. This allows us to work out the terms of the covariance matrix relating **T** and **F**_mean_.

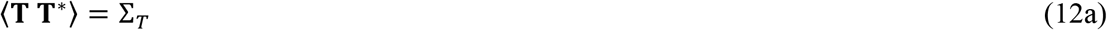

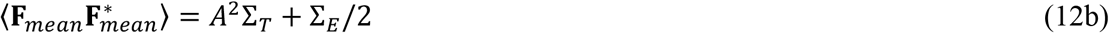

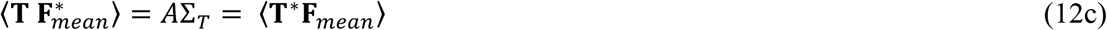

The likelihood target has a simpler form if it is expressed in terms of normalised map coefficients (E-values), in which the mean-square value of E is expected to be one.

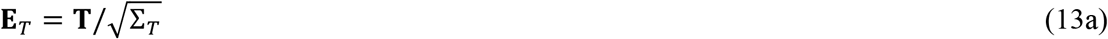

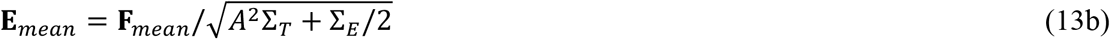

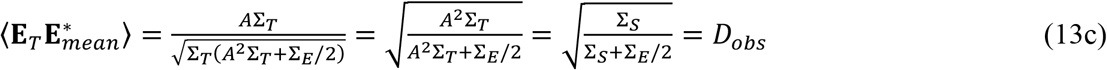

*D_obs_* is the complex correlation relating **E**_*mean*_ and the true value, **E**_*T*_. It plays the same role, for a single Fourier term, as *FSC_ref_* (Fourier shell correlation to the true map) does for a whole resolution shell in Fourier space. Note that, if Σ_*E*_ is zero, *D_obs_* is one, but it becomes smaller as the ratio between Σ_*E*_ and Σ_*S*_ increases, reaching zero when Σ_*S*_ is zero.

The other source of error in the likelihood target is model error. For docking, it is generally safe to assume that the errors in the map and the errors in the model are independent, prior to any refinement against the map, so there are no concerns about overfitting. The relationship between the Fourier coefficients computed from a model and those that would be obtained from the true map is the same as that between calculated and true structure factors in crystallography: the Central Limit Theorem allows us to conclude that the errors in the calculated Fourier coefficients that arise from the sum of many small errors from the individual atoms in the model can be described in terms of a complex normal distribution, like the errors between the true map and the experimental reconstruction. In crystallography, this is described by the complex correlation, termed *σ_A_*, between the normalised structure factors (Srinivasan & Ramachandran, 1965; Read, 1990). The *σ_A_* term combines the effects of completeness of the model (the fraction *f* of the scattering accounted for by the model) and the accuracy of the model; if we make the simplifying assumption that the errors in the coordinates of all the atoms are all drawn from the same 3D Gaussian distribution, *σ_A_* can be calculated with the following formula:

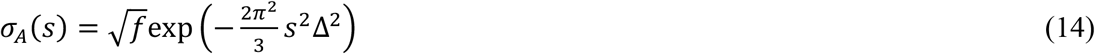

where *s* is the inverse resolution and Δ is the rms radial coordinate error (Read, 1990). Violation of this assumption can change the resolution dependence of the *σ_A_* curve, but a compromise effective overall rms error is determined by refinement after placing the model. The complex normal distribution is relevant regardless of the types of modelling errors, as long as none of them are so large that they dominate.

Because the errors between the true map and either the calculated map or the observed map are independent, the complex correlation between the observed map and the model is simply the product of the two individual complex correlations, *D_obs_* and *σ_A_*. Therefore, the joint distribution of **E**_*mean*_ and the normalised calculated Fourier coefficient, **E**_*C*_, is a bivariate complex normal distribution with expected values of zero (prior to any knowledge of either) and the following covariance matrix:

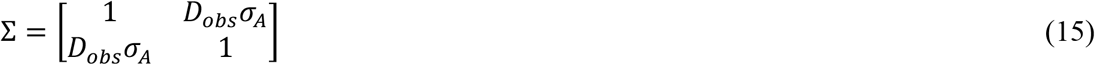

The likelihood function for judging the fit of a model in a map (whether derived by docking or structure refinement) is the conditional probability distribution for the observed normalised Fourier coefficient given the corresponding term computed from the model. This conditional distribution is obtained by a simple manipulation of the joint distribution, yielding a complex normal distribution with a variance of 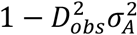 and an expected value of *D_obs_σ_A_***E**_*C*_:

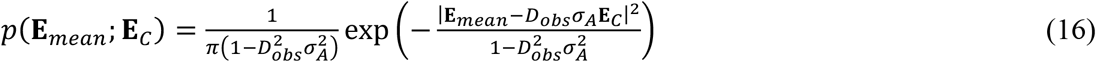

It is more convenient to work with the log-likelihood gain or LLG, *i.e*. the gain in the log-likelihood score compared to an uninformative model (for which *σ_A_* is zero). The contribution of a single Fourier term to the total LLG can be determined by taking the logarithm of *p*(**E**_*mean*_; **E**_*C*_) and subtracting the logarithm of that probability with *σ_A_* set to zero. After expanding the arguments of the exponentials in terms of the amplitudes and phases of the Fourier terms, the result can be simplified to the following:

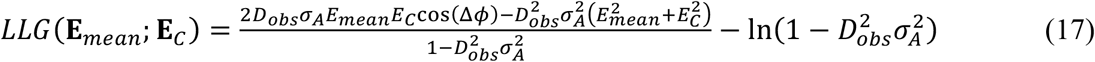

where Δ*ϕ* is the difference between the phases of **E**_*mean*_ and **E**_*C*_, and **E**_*mean*_ and **E**_*C*_ are amplitudes. Note that this can alternatively be expressed in terms of a correlation function between the weighted averaged map and the model, a scale factor and an offset:

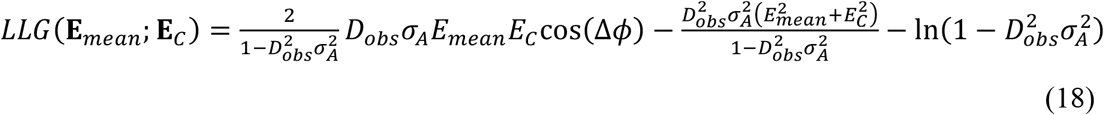

The total LLG score is the sum over all Fourier terms. However, it is important to note that cryo-EM differs from crystallography in that the Fourier transform of the reconstruction is typically highly over-sampled: proteins in crystals pack in a lattice where molecules are in contact with each other, whereas the cryo-EM reconstruction is computed in a much larger box than required to contain the particle. Oversampling leads to strong correlations among neighbouring Fourier terms. This can be accounted for simply by applying a correction factor equal to the ratio of the volume required to contain the particle and the volume of the box in which the reconstruction was carried out, as proposed by van Heel and Schatz (van Heel & Schatz, 2020) for computing information content. The same correction factor must be applied to all fast approximations, expected values and information gain discussed below.

The likelihood target described here has the same basic functional form as the likelihood target used to refine cryo-EM models in Refmac (Murshudov, 2016), differing importantly in taking account of the dependence of the signal and error terms on direction in reciprocal space, instead of depending only on resolution. As shown in section 5 below, we see large differences in the size of signal and error terms within a resolution shell, which will have significant effects on the likelihood scores.

### 2.3. Map coefficients

Two sets of map coefficients can be generated for evaluating the fit of a docked model. The first type uses the Fourier coefficients

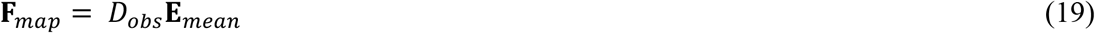

Since this is the expected value of the true sharpened map coefficient (the centroid of the probability distribution), this should give a map that minimises the error from the true sharpened map.

The second type uses Fourier coefficients that include the other weighting terms from the correlation function in the log-likelihood target (18),

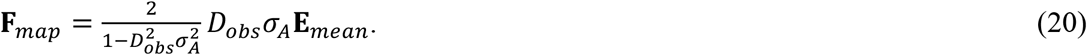

The correlation of this map (20) with the sharpened map computed from a docked model should be proportional to the likelihood target. To compute such a map, a choice has to be made for the value of *σ_A_* that is used, which primarily depends on the scattering in the volume under consideration but also coordinate error and on the ability of atomic models to account for the bulk solvent region.

### 2.4. Fast rotation target for scoring orientations of models

In crystallographic MR, the six-dimensional problem of finding the orientation and position of a model to fit the diffraction data is typically divided into a sequence of two threedimensional problems: an orientation search (rotation function) followed by translation searches with models in a number of plausible orientations determined from the orientation search (translation function). The crystallographic rotation function can be directly adapted to the docking problem in cryo-EM; it does not use the phase information in the complex Fourier terms, but phase information cannot be used in any event without some knowledge of the position of the search model. The rotation search thus depends solely on the amplitudes of the Fourier terms, so the crystallographic likelihood-based rotation function (Storoni *et al*., 2004) can be used without alteration: the cryo-EM *D_obs_* plays the same role as the crystallographic *D_obs_* parameter in the log-likelihood-gain on intensities (LLGI) target (Read & McCoy, 2016; Jamshidiha *et al*., 2019). As in crystallography, an approximation of the likelihood-based rotation function can be computed rapidly by FFT methods before being scored by the exact likelihood function.

It should be noted that phase information can be used indirectly in the rotation search. If there is a hypothesis for the location of a particular component in a full reconstruction, the rotation search can use the Fourier terms computed from a portion extracted from the full reconstruction. The use of such a procedure is essential to the sub-volume searches mentioned below and discussed in detail in the accompanying paper (Millan *et al*., 2023).

### 2.5. Fast translation target for scoring positions of oriented models

In crystallography, where there is typically no prior phase information in an MR search, only an approximation to the likelihood target can be computed by FFT methods. However, the LLG score for the fit of a model to cryo-EM data (18) takes the form of a correlation function, which can therefore be calculated exactly as a function of translation using a single FFT, plus terms accounting for scaling parameters and contributions that do not change with translation.

### 2.6. Rigid-body refinement

Refinement of a docked model involves optimizing parameters of (18) to maximise the LLG. The orientation and translation parameters affect the calculated Fourier terms, while the estimated RMSD of the model from the true structure after being correctly placed (typically in the range of 0.8-1.2 Å) changes the *σ_A_* term. As in the related MR case, a careful choice of parameterization can improve the refinement behaviour. For instance, correlations between rotation and translation parameters can be minimised by defining the rotation in terms of a rotation about the center of mass of the component. In addition, defining the rotation in terms of a perturbation applied to the current orientation by rotating sequentially about orthogonal x, y and z axes (rather than, for instance, Euler angles) makes the rotation parameters locally close to orthogonal.

Although improvements in hardware and cryo-EM protocols have generally reduced the uncertainty about voxel size (or magnification factor) in modern cryo-EM reconstructions, we have implemented a cell scale factor parameter, which affects the calculated Fourier terms and therefore can be refined to compensate for any error in voxel size.

## 3. Expected likelihood scores and information gain

In crystallographic MR, it has been possible to optimise the choice of search strategy by, first, knowing what absolute LLG score is required for correct solutions to be recognised and, second, being able to predict the LLG score that can be achieved in a particular search, given the quality of the model, the quality of the data and the resolution limit applied to the data (McCoy *et al*., 2017; Oeffner *et al*., 2018). The same considerations of expected LLG (or eLLG) can be applied to docking in cryo-EM, as discussed in detail in the accompanying paper (Millan *et al*., 2023).

### 3.1. Rotation eLLG

In a rotation search for a cryo-EM reconstruction lacking symmetry, the LLG score for an orientation is the same as the crystallographic LLG score for a model of a crystal in space group P1. Therefore, the rotation eLLG, *eLLG_rot_*, can be computed with the same formula as the crystallographic eLLG for space group P1 (McCoy *et al*., 2017):

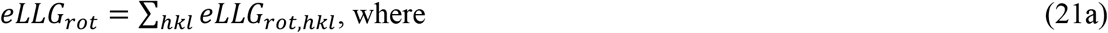

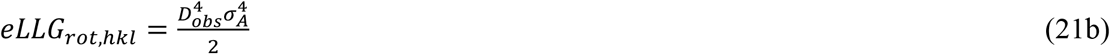

In (21), the value of *D_obs_* for each Fourier term is obtained from the analysis of the reconstruction or a sub-volume extracted from it. It is instructive to consider the effect of increasing the volume of a sphere extracted from the total reconstruction. If a sphere containing the correct volume for the component under investigation were doubled in volume, the number of Fourier terms would double. At the same time, the fraction of the map accounted for by the model would decrease by the same factor of two. Because *σ_A_* is proportional to the square root of the model completeness, each term in the sum would be reduced by a factor of 4, so that the total *eLLG_rot_* would be reduced by a factor of two. More generally, all else being equal, *eLLG_rot_* is inversely proportional to the volume of the part of the map being used for the search.

### 3.2. Translation eLLG

The expected value of the LLG for an individual Fourier term is given by the probability-weighted average of the LLG over all possible values of the calculated Fourier term, where the weighting is the conditional probability of that calculated term given the observed Fourier term. Because the joint probability distribution of the calculated and observed Fourier terms is symmetric, the required conditional probability has the same functional form as the likelihood of the data given the model.

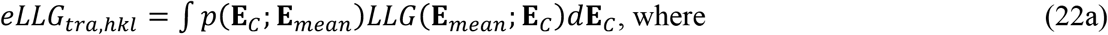

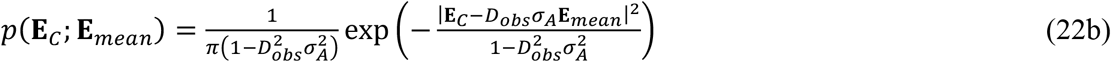

The integral has a simple analytical solution, which was determined using *Mathematica* (Wolfram Research; Inc., 2019):

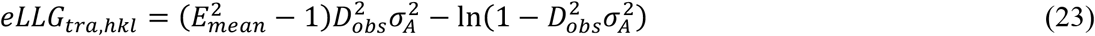

Considering that the expected value of 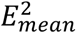 is one, if we assume that there is no correlation between 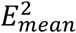 and 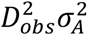, the expected value of the first term is zero, so that

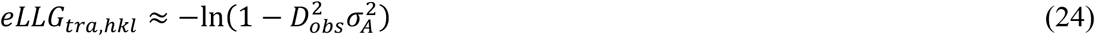

For all values of *D_obs_σ_A_, eLLG_tra,hkl_* is greater than *eLLG_rot,hkl_*, especially for the poorest combinations of map and model quality; when *D_obs_σ_A_* is 0.01, for instance, the ratio is about 20,000. This is an indication of the extent to which phase information enhances the likelihood scores. The implication is that the trade-off between the size of the sub-volume and the sensitivity to the correct solution is very different for the rotation and translation parts of the search.

In contrast to *eLLG_rot,hkl_*, *eLLG_tra,hkl_* is relatively insensitive to the size of sub-volume, especially when either the map or the model is poor (*D_obs_σ_A_* « 1), in which case the change in number of Fourier terms counterbalances the change in the logarithmic term. Intuitively this makes sense, because the inclusion of phase information implicitly focuses the calculation on the position of the model. The implication that the size of the sub-volume is relatively unimportant for the translation search can be exploited for a 6-dimensional bruteforce search, as discussed in the accompanying paper (Millan *et al*., 2023).

### 3.3. Information gained by cryo-EM reconstruction

Information theory and likelihood are closely connected, and the information gained by measuring the data in a cryo-EM reconstruction, computed using the Kullback-Leibler divergence (Kullback & Leibler, 1951) can be derived with methods related to those used for the eLLG. Essentially, the Kullback-Leibler divergence (if measured with the natural logarithm in units of natural units of information, nats, rather than the conventional bits obtained with the logarithm base 2) is equivalent to the eLLG that would be expected for a perfect model (*i.e*. the true structure in the sample).

The Kullback-Leibler divergence for the information gained about the true map, given the reconstruction, can be computed for one Fourier term with the following integral over all possible values of the true Fourier term, **E**:

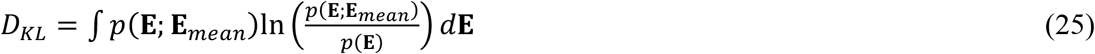

The probability weighting this integral is equivalent to *p*(**E**_*C*_; **E**_*mean*_) when the model is perfect (i.e. **E**_*C*_ = **E**). Applying Bayes’ theorem, we can substitute

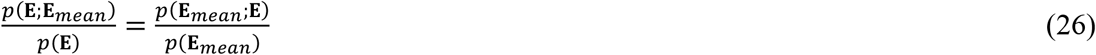

for the likelihood ratio in the logarithm. The logarithm of a likelihood ratio is equivalent to the differences between the logarithms of the likelihoods, so we see that the logarithm in the integral is the LLG that would be achieved with a perfect model. The expression for *D_KL_* is therefore equivalent to the expression for *eLLG_trahkl_* in (22a) if the model were perfect. Because a perfect model would have *σ_A_* = 1 for all Fourier terms,

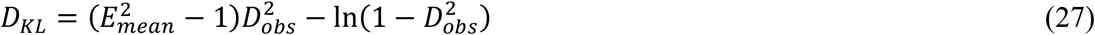

Noting as before that the mean value of 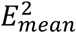 should be one, if 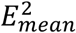 and 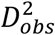 are uncorrelated we have

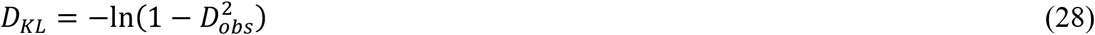

Information in units of bits instead of nats can be obtained by using the logarithm base 2, which differs by a factor of ln(2).

The total information gain in an entire data set or in a resolution shell will be the sum from the individual Fourier terms, but corrected for the correlations arising from oversampling in Fourier space. As above, following similar reasoning to that invoked by van Heel and Schatz (2020), the correction for oversampling can be made by comparing the volume of the map to the volume occupied by the ordered part from which the signal is obtained.

Although it is not immediately obvious, the *D_KL_* measure proposed here is closely related to the information content measure proposed by van Heel and Schatz (2020), in which they followed a different line of reasoning. They proposed a Fourier Shell Information (*FSI* measure for the information, in bits, gained by a shell of data in Fourier space, expressed in terms of the *FSC* between two half-maps for that resolution shell:

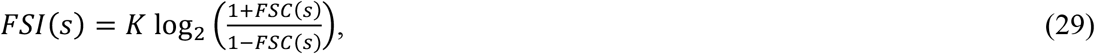

where *K* is the effective number of independent Fourier terms in the shell under consideration. As noted above, *D_obs_* plays the same role for a single Fourier term as *FSC_re_f* does for a resolution shell. If we assume that all Fourier terms in a resolution shell have the same value of *D_obs_*, we can express the *FSI* equation in terms of *D_obs_* using the relationship between *FSC* and *FSC_ref_* derived by Rosenthal and Henderson (2003).

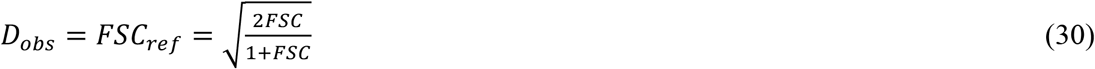

Solving for *FSC* yields

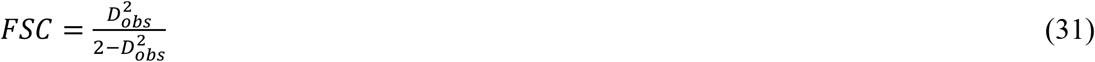

Substituting this for FSC in (29) and simplifying yields the following:

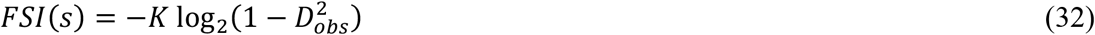

Interpreting *K* as the number of independent Fourier terms in the shell, this is equivalent to the expression given above for the Kullback-Leibler divergence measured in bits. The expression given here is more general, because it allows for differences in accuracy of different Fourier terms around a shell, arising from anisotropy and the effects of favoured orientations, which will lead to variation among the values of *D_obs_* for different terms.

In our docking calculations, the information gain calculation is used to save computing time by omitting Fourier terms that will have almost no effect on the likelihood calculation. As done in the related molecular replacement calculation (Jamshidiha *et al*., 2019), Fourier terms with an information gain of less than 0.01 bit are ignored after the error analysis step.

## 4. Implementation of algorithms

The algorithms have been implemented as a combination of Python scripts and C++ code, both making substantial use of the Computational Crystallography Toolbox, cctbx (Grosse-Kunstleve *et al*., 2002).

Tools to analyze the maps, determine the parameters characterizing the signal and noise, and compute modified Fourier coefficients for the docking calculation have been implemented in the Python program *prepare_map_Jor_dockmg*.

The *prepare_ map_for_ docking* tool is available as a Python script within the *maptbx* section of the open-source Computational Crystallography Toolbox, *cctbx* (Grosse-Kunstleve *et al*., 2002). This is available standalone and also as part of the *Phenix* (Liebschner *et al*., 2019) and *CCP4* (Winn *et al*., 2011) software suites.

## 5. Results

### 5.1. Behaviour of signal and error analysis

As noted by Palmer and Aylett (2022), errors are similar throughout a cryo-EM reconstruction, but signal-to-noise ratios can vary dramatically within the reconstruction because of variations in the strength of the signal. This can be demonstrated by looking at the local behaviour of the signal power (Σ_*S*_) and noise power (Σ_*E*_) in reciprocal space, after the analysis using the *prepare_map_for docking* tool. One informative example is the map for conformation 2 of the *E. coli* respiratory complex I (EMDB entry 12654, PDB entry 7nyu), for which the local reconstruction quality varies widely (Kolata & Efremov, 2021). An analysis of one of the best and one of the worst regions of the map is given in Fig. 1, illustrating that the noise power is similar in the two regions whereas the signal power, and its variation in Fourier space, differs substantially. The poorly-ordered region of the map corresponds to chain L of the model, where the authors estimate the local resolution to be in the range of 9-11 Å. This agrees roughly with the resolution at which the signal and noise powers are equivalent, in Fig. 1b.

**Figure 1.**
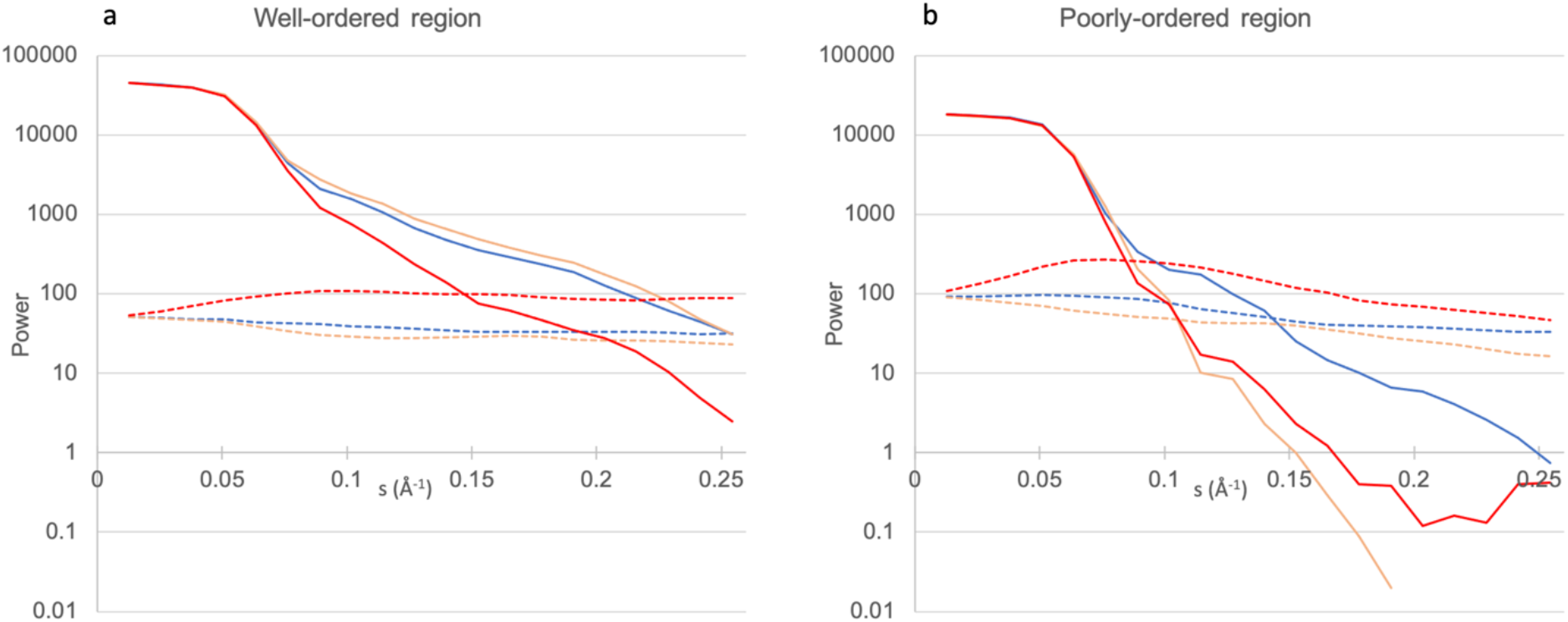
Variation of signal power (Σ_*S*_) and noise power (Σ_*E*_) in Fourier space for a) well-ordered (center of cytoplasmic domain) and b) poorly-ordered (chain L) regions of the *E. coli* respiratory complex I. Both regions correspond to spheres with a radius of 30 Å. Signal power is shown with solid lines and noise power with dashed lines, as it varies in three directions parallel to the x (blue), y (orange) and z (red) coordinate axes of the reconstruction. The variation of noise power is similar for the two regions of the reconstructions, but the variation of signal power differs significantly, even in the relative falloff in the three directions.

## 6. Discussion and conclusions

The problem of docking an atomic model into a cryo-EM reconstruction is reminiscent of the molecular replacement problem in crystallography. The similarity is more than superficial, as both problems can be addressed using likelihood functions that start from joint distributions of complex Fourier terms. In both cases, the model is represented by its Fourier transform (either of its electrostatic potential or its electron density), but cryo-EM differs in the important fact that the data retain the phase information lost in the crystallographic diffraction experiment.

Applying likelihood requires characterising all sources of error, which differ between the methods. In cryo-EM, the typical presence of favoured particle orientations leads to large differences in the reliability of the Fourier terms. The variation of noise contributions to the Fourier terms is expected to vary smoothly over Fourier space, and a method to assess this variation has been developed.

The likelihood framework allows the implementation of tools that have been found useful in molecular replacement. In particular the expected log-likelihood-gain (eLLG) score can be calculated in advance of any docking search, as well as the information gained by making the cryo-EM reconstruction.

The accompanying paper (Millan *et al*., 2023) describes the implementation of these ideas in software tools for docking, and the success of those tools demonstrates the validity of the approach described here, including the use of eLLG to choose optimal strategies.

## Funding information

This research was supported by the Wellcome Trust (grant 209407/Z/17/Z to RJR) and the National Institutes of Health (grant GM063210 to TCT and RJR).

## Notes

### Competing Interest Statement

The authors have declared no competing interest.

### Summary of Updates

Revised in response to referee comments to clarify explanations.

